# PDZ-containing proteins targeted by the ACE2 receptor

**DOI:** 10.1101/2021.07.23.453470

**Authors:** Célia Caillet-Saguy, Nicolas Wolff

## Abstract

Angiotensin converting enzyme 2 (ACE2) is a main receptor for SARS-CoV-2 entry to the host cell. Indeed, the first step in viral entry is the binding of the viral trimeric spike protein to ACE2. Abundantly present in human epithelial cells of many organs, ACE2 is also expressed in the human brain. ACE2 is a type I membrane protein with an extracellular N-terminal peptidase domain and a C-terminal collectrin-like domain that ends with a single transmembrane helix and an intracellular 44-residues segment. This C-terminal segment contains a PDZ-binding motif (PBM) targeting protein interacting domains called PSD-95/Dlg/ZO-1 (PDZ).

Here, we identified the human PDZ specificity profile of the ACE2 PBM using the high throughput holdup assay and measuring the binding intensities of the PBM of ACE2 against the full human PDZome. We discovered 14 human PDZ binders of ACE2 showing significant binding with dissociation constants values ranging from 3 to 81 μM. NHERF, SHANK, and SNX27 proteins found in this study are involved in protein trafficking. The PDZ/PBM interactions with ACE2 could play a role on ACE2 internalization and recycling that could benefit for the virus entry. Interestingly, most of the ACE2 partners we identified are expressed in neuronal cells, such as SHANK and MAST families, and modifications of the interactions between ACE2 and these neuronal proteins may be involved in neurological symptoms of COVID-19.

## Introduction

Three human coronaviruses have appeared since the early 2000s and caused epidemics: SARS-CoV (2003), MERS-CoV (2012), and finally SARS-CoV-2 (2019). However, only SARS-CoV-2 has become pandemic in a few months.

SARS-CoV-2 is a 30-kb RNA virus that belongs to the *betacoronavirus* genus and must enter a host cell in order to be able to replicate. The first step in this process is therefore the entry of viral material into the cytoplasm after passing through the cell membrane. The entry step begins with the attachment of the viral particle to the cell surface. This is based on the interaction between the spicules on the surface of the viral particle (spike protein S of SARS-CoV-2) and the glycoprotein angiotensin-converting enzyme 2 (ACE2) which acts as an entry receptor [1].

After attachment to ACE2, the viral protein S is cut into two parts (S1 and S2) by the serine protease TMPRSS2 which is present on the surface of the host cell [2]. Then the furin cleaves the S2 region (S2’) to start conformational changes for fusion between the viral envelope and the plasma membrane of the host cell. The “fusion peptide”, part of the polypeptide sequence of S, will be exposed, which fits into the cell membrane. There follows a gathering between the envelope of the virus and the cell membrane, both formed by a lipid bilayer which will therefore fuse [3]. In a second ACE2-dependent route, the virus can also enter by endocytosis by the cathepsin-L pathway. Cathepsin L is ubiquitously expressed in mammalian cells and cleaves the S2′ and activates the fusion between viral and endosomal membranes, leading to the release of the viral RNA into the host cell cytoplasm [4] where the virus replicates [5].

The presence of the ACE2 viral receptor is a major determinant of the specific recognition between the virus and the host. SARS-CoV-2 can therefore infect human cells expressing ACE2. The expression profile of ACE2 is broad. The highest expression levels of ACE2 are found in the small intestine, testis, kidneys, heart, thyroid and adipose tissue; intermediate in the lungs, colon, liver, bladder and adrenal glands; lowest in the blood, spleen, bone marrow, brain, blood vessels and muscle [6]. Symptomatic patients with SARS-CoV-2 infection are often reported having fever, cough, nasal congestion, fatigue, and other signs of an upper respiratory tract infection, which can develop into acute respiratory distress syndrome (ARDS) with a low survival rate [7]. Furthermore, ACE2 is essential to the regulation of the renin-angiotensin-aldosterone system (RAAS), which is a main regulator of blood pressure as well as fluid and electrolyte homeostasis.

ACE2 is a glycosylated integral membrane protein of 805 residues with a large N-terminal ectodomain (residues 18-740), a transmembrane domain of 21 residues (residues 741-761) and a short cytoplasmic C-terminal region (residues 762-805) containing a PDZ-binding motif (PBM) at the C-terminal extremity (residues 803-805) [8](Figure 1A). This PBM of class I with the motif sequence X-(S/T)-X-Φ_COOH_, where X is any residue (position -1), and Φ is a hydrophobic residue (last position 0)(Figure 1B), allows to bind to other proteins through their PDZ (PSD-95/Dlg/ZO-1) domains. The affinities of a FITC-labeled C-terminal peptide of ACE2_796–805_ were previously measured for 8 PDZ domains by fluorescence polarization assays [9]. Three PDZ domains, the first PDZ domain (PDZ1) of NHERF3 and the PDZ domains of SHANK1 and SNX27, were found to bind to the ACE2 PBM peptide. However, only 8 over 272 PDZ expressed in the human proteome (the PDZome) have been tested in this study. Recently, ACE2 was also shown to interact with NHERF1 PDZ domains enhancing ACE2 membrane residence, and consequently increasing the ACE2-mediated SARS-CoV2 cell entry [10].

**Figure 1:**
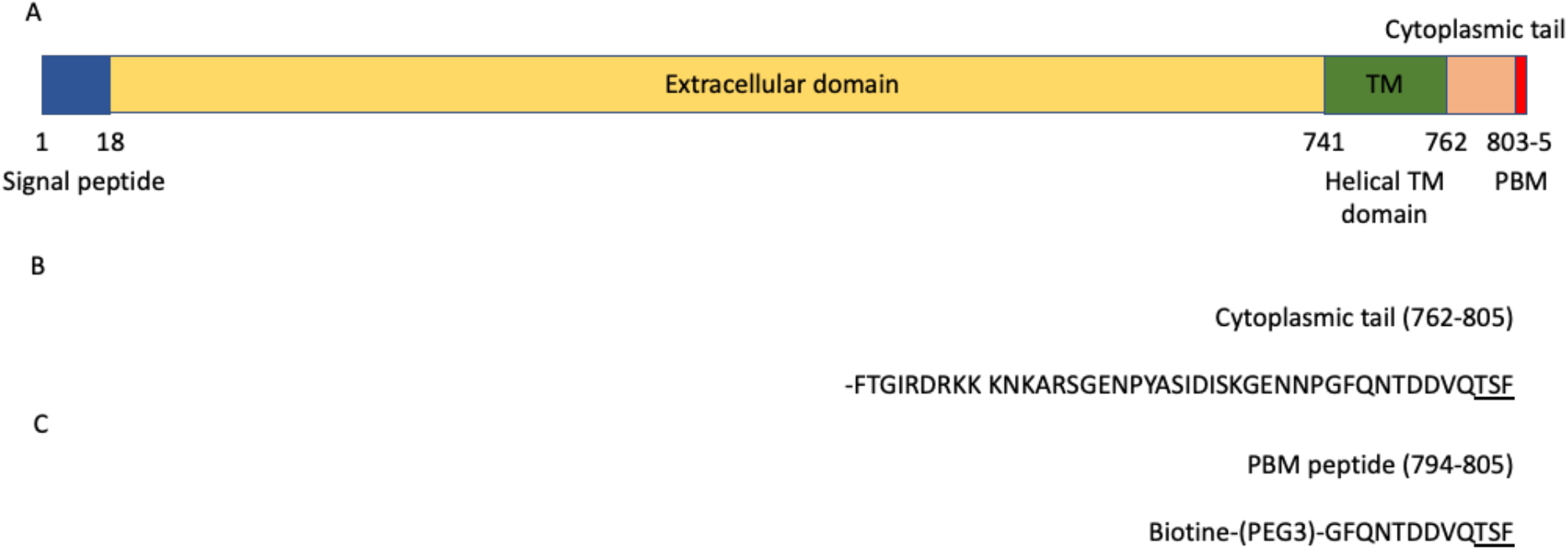
ACE2 domain organization. A) Schematic domain organization of the ACE2 protein. B) Sequence of the cytoplasmic tail (762-805) of ACE2. C) Sequence of the peptide used in holdup assay. The PBM is underlined in the sequences.

The PBM-PDZ interactions are involved in a wide variety of critical cellular processes and are often disturbed in pathologies and by viral proteins during infection. The human proteome contains 272 PDZ domains distributed over 152 proteins. Recently, we used the holdup assay [11] adapted to low–to-medium (1-100 μM) affinity, for high-throughput quantification of PDZ-PBM interaction to screen the PBMs of SARS-CoV-2 E, 3a and N proteins against the PDZome [12].

In this study, we used the holdup assay to identify the cellular PDZ-containing proteins targeted by ACE2 through a PDZ-PBM interaction among the full PDZome. We establish a list of PDZ-containing partners potentially involved in ACE2 cell signaling through its PBM. We identified 14 PDZs that significantly bind to the ACE2 PBM and determined their affinities. Interestingly, NHERF, SHANK, and SNX27 proteins found in this study could play a role on ACE2 internalization and recycling that could benefit for the virus entry. In addition, the ACE2 PDZ-binding motif binds notably to neuronal proteins, SHANK and MAST1/MAST2 proteins, that could participate to the neurological alterations in infected patients by SARS-CoV-2.

## Results

### The ACE2 PDZ-binding motif recognizes 14 PDZ domains with affinities values of interactions ranging from 3 μM to 81 μM

To investigate the specificity profile of the PBM of ACE2 against all human PDZ domains, we used the automated high-throughput holdup assay, which allows measuring binding intensities (BIs) for a large number of PDZ domain-PBM pairs [11]. We used an updated PDZome library that contains all the 266 human PDZ domains [13]. The ACE2 PBM peptide was synthetized as a 12-mer encompassing the C-terminal PBM sequence of ACE2 protein linked to a N-terminal biotinyl group (Figure 1C). The PDZome-binding profile (Figure 2A) represents the individual BIs of each PDZ domain for the ACE2 PBM peptide. The binding intensities are directly linked to the PBM/PDZ affinities. Thus, the holdup approach shows a high sensitivity to detect low–to-medium affinity PDZ/PBM pairs. It provides an affinity-based ranking of the identified binders corresponding to a specificity profile (figure 2A). Based on BIs values higher than 0.2 for significant PDZ-peptide interactions [11], 14 PDZ domains were identified (Figure 2B, Table 1). This dataset represents about 5% of the human PDZome targeted by the PBM of ACE2 as illustrated by the sharp profile of interaction of Fig. 2A reflecting the specificity of these interactions.

**Figure 2:**
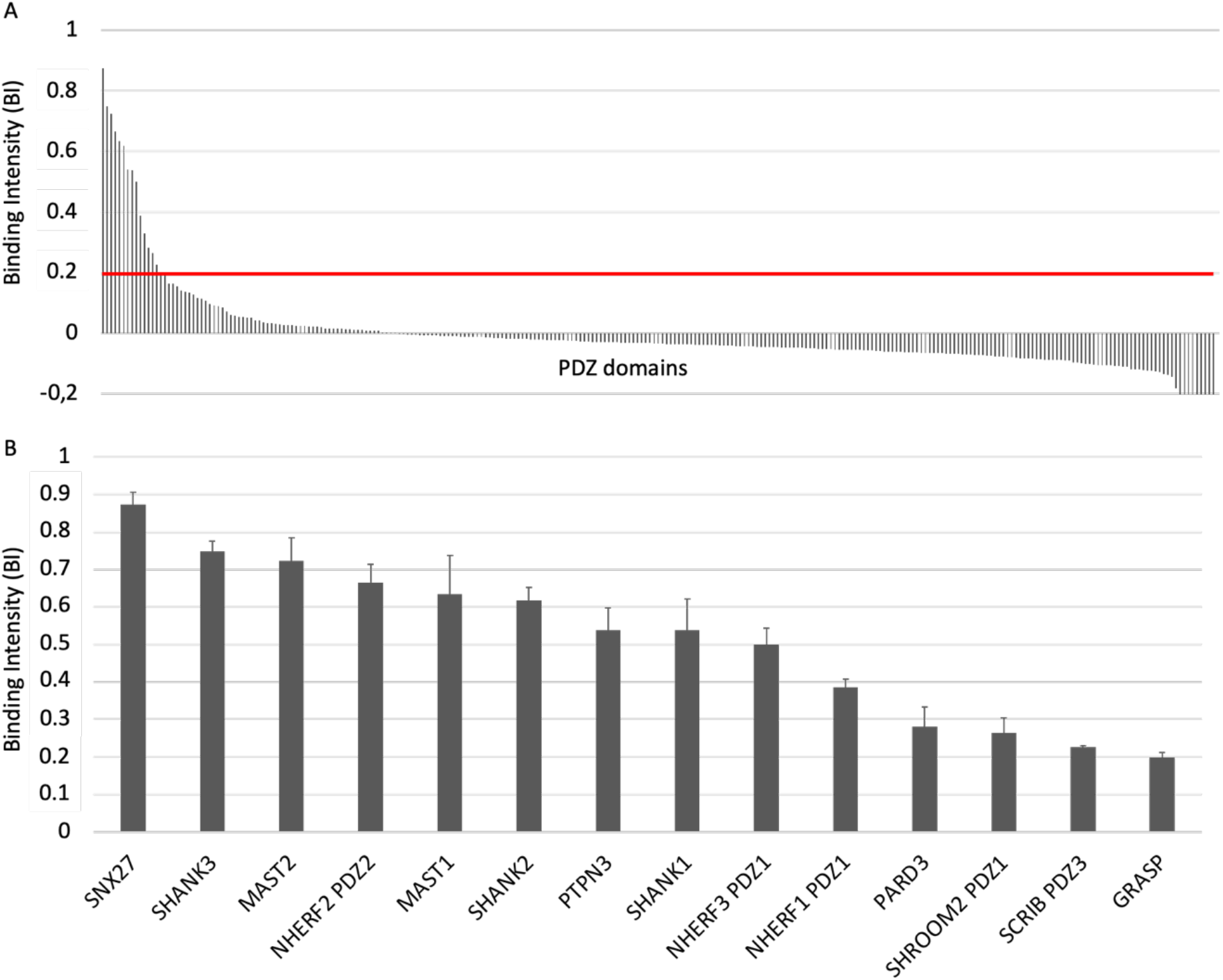
PDZome-binding profiles of ACE2 protein PBM by holdup assay. A) PDZ domains are ranked by decreased BI values. A red line indicates the threshold of significant BI at 0.2. B) Zoomed-in view shows the 14 PDZ binders of ACE2 PBM peptide, which displayed BI values higher than 0.2. The data are representative of two independent experiments and error bars correspond to the standard deviation.

We convert the holdup BI values to Kd values as described previously [14,15]. Using the minimal threshold BI value of 0.2 corresponding to a Kd value of ∼80 μM dissociation constant in this study, the fourteen PDZ binders of ACE2 PBM showed interactions with affinities values ranging from 3 μM (SNX27) to 81 μM (GRASP) (Fig. 2B, Table 1). Thus, the canonical PBM sequence of ACE2 protein binds *in vitro* to several PDZ domains in the 1-100 μM affinity range typically found for PDZ/PBM interactions. The PDZ of SNX27 is the best binder of ACE2 PBM in our assay, while three other PDZs present Kd values in the micromolar range: the PDZ domains of SHANK3 and of MAST2, and PDZ2 of NHERF2 (Table 1). These are considered as strong interactions for PDZ/PBM complexes. Then, the PDZ domains of MAST1, SHANK1, SHANK2, PTPN3, PARD3, GRASP, PDZ1 of NHERF3 and NHERF1, of SHROOM2, and the PDZ3 of SCRIB have intermediate Kd values between 10 and 81 μM for the ACE2 PBM peptide.

### The PDZ-binding motif of ACE2 targets proteins involved in protein trafficking and some proteins preferentially expressed in neurons

SNX27 is the best binder of ACE2 PBM in our holdup assay. This binding was previously reported by Kliche *et al*. [9]. SNX27 belongs to the sorting nexin family of proteins and is involved in the recycling of internalized transmembrane proteins [16]. SNX27 is widely expressed but has been extensively studied in the context of the central nervous system (CNS) homeostasis regulation [17]. SNX27 is important for higher-order processes, such as learning and memory [18] and patients with SNX27 variants display seizures, developmental delay, behavioral disturbance, and subcortical brain abnormalities [19].

SHANK1, SHANK2 and SHANK3 are targeted by the ACE2 PBM and belong to the SHANK family composed of developmental genes involved in the development and function of neural circuits, as well as in the formation of synapses. Mutations in SHANK genes involve autism spectrum disorder with altered functions in synaptic scaffolding to regulating spine morphology and neurotransmission [20].

The three members of the NHERF family (Na+/H+ exchanger regulatory factor also called EBP50), NHERF1, NHERF2 and NHERF3, are also targeted by the PBM of ACE2. The NHERF family functions are broad [21]. These scaffold proteins contain two PDZ domains and a C-terminal sequence that binds several members of the ERM (ezrin-radixin-moesin) family and they are highly expressed in epithelial tissues [22]. Many NHERF partners have been identified broadening the protein functions in particular in membrane physiology. NHERF is involved in transport proteins to apical membranes and in regulation of endocytosis and internal trafficking of membrane proteins [23]. NHERF1 was recently shown to interact through PDZ/PBM interaction with ACE2. Zhang *et al* proposed that this interaction facilitates ACE2 receptor internalization and thus SARS-CoV-2 virion internalization since it is attached through ACE2 by the viral surface spike protein [10].

Then, two members of the MAST family, MAST1 and MAST2, are targeted by the PBM of ACE2. The MAST family is composed of five genes (MAST1, MAST2, MAST3, MAST4, and MASTL) and is predominantly expressed in neuronal cells [24]. It forms a subgroup of microtubule associated serine/threonine kinases composed of a kinase domain and a PDZ domain. They are involved in microtubule assembly, an essential step in neurogenesis. MAST1 is essential for correct brain development [25] and MAST1 substitutions are present in patients with autism and microcephaly, suggesting that mutations in this gene might be involved in neurodevelopmental diseases. MAST2 is an anti-survival kinase that interacts with PTEN in neurons [26] among other roles. MAST2 shows association with red blood cell distribution width which was identified recently as a biomarker of COVID19 mortality [27]. MAST3 may play a role in developmental and epileptic encephalopathies [28].

PTPN3 is a tyrosine phosphatase, the vertebrate homolog of Ptpmeg, expressed among others in the nervous system and involved in neuronal circuit formation in the drosophila central brain [29]. PTPN3 is also often found involved in a variety of human cancers [30].

PARD3 is essential in epithelial tight junctions. It is required for establishment of neuronal polarity and normal axon formation in cultured hippocampal *neurons* [31,32]. Interestingly, we previously showed that is a potential target of the SARS-CoV-2 envelope PBM [12]. SCRIB is a polarity protein important in maintaining cell junctions and belongs to polarity complexes as PARD3.

SHROOM2 may be involved in endothelial cell morphology changes during cell spreading [33] and GRASP (general receptor for phosphoinositides 1-associated scaffold protein also named Tamalin) regulates the trafficking of metabotropic glutamate receptors that play important roles in various neuronal functions [34].

Hence, 9 over 14 proteins have been described to play a role in neuronal cells depicting a neuronal-oriented specificity profile with a significant emphasis on cellular junctions and trafficking.

## Discussion

ACE2 is involved in blood pressure regulation and cleaves a range of substrates involved in different physiological processes. ACE2 receptors and proteases such as transmembrane serine protease 2 (TMPRSS2) and furin are important for viral entry into host cells for coronaviruses such as SARS-CoV-2. Importantly, it is the functional receptor for SARS-CoV-2 responsible of the COVID-19 disease. ACE2 belongs to a family of transmembrane proteins that has wide tissue distribution. Protein expression of ACE2 was initially identified in the heart, kidney, and testis, on the surface of cells that are in contact with the cellular environment like the lung’s alveolar epithelial cells and the small intestine’s enterocytes. In addition, ACE2 is also localized in the brain [35,36].

In this work, we demonstrate that a peptide mimicking the C-terminal sequence of ACE2 containing a 3-residues canonical class I PBM binds in vitro to several human PDZ domains. Fourteen PDZ domains (5% of the human PDZome) were identified as binders of the PBM of ACE2 with Kd values ranging from 3 μM to 81 μM.

The 14 PDZ-PBM interactions found in this study may impact the SARS-CoV-2 viral infection as recently observed for NHERF1 and SNX27. In this case, the PDZ/PBM interaction of ACE2 with NHERF1 facilitates SARS-CoV-2 internalization by enhancing the ACE2 membrane residence [10]. Interestingly, the four PDZ-containing proteins, PARD3, NHERF1, MAST2 and SNX27, were also identified as binders of the PBM of E protein of SARS-CoV-2 in our previous study [12]. Among these binders, we showed that PARD3 and MAST2 significantly affect viral replication under knock down gene expression in infected cells. This suggests that SARS-CoV-2 also targets ACE2 partners to potentially facilitate virus entry. NHERF, SHANK, and SNX27 proteins are involved in receptor trafficking both at the cell surface and intracellular side that could be linked to the recycling of ACE2. SNX27 is involved in transport by the recycling of internalized transmembrane proteins [37]. It is required for SARS-CoV-2 entry [38]. Thus, SNX27 might be involved in recycling ACE2 to the plasma membrane stimulating viral entry as previously suggested by Kliche *et al*. [9]. The NHERF family is also a potential target of the ACE2 PBM. NHERF2 is structurally related to NHERF1. NHERF3 regulates the surface expression of plasma membrane proteins in the apical area of epithelial cells in agreement with the enrichment of ACE2 on the apical surface of airway epithelia [39]. Thus, PDZ/PBM interactions can play a role on ACE2 localization and recycling, which could be gainful for the virus in particular for internalization.

We found that nine PDZ-containing proteins targeted by ACE2 PBM display neuronal functions: the three SHANK, two MAST proteins, SNX27, PARD3, GRASP and PTPN3. Interestingly, the human PDZ specificity profiles we previously established for neurotropic viruses with the PBMs of the Rabies glycoprotein and of the West Nile NS5 protein mainly overlap the PDZ profile we showed in this study [40,41]. ACE2 in the CNS is involved in the regulation of cardiovascular function [42] but not only. ACE2 has noticeable effects on blood pressure, cardiac hypertrophy, stress response, anxiety, cognition, brain injury and neurogenesis [43]. ACE2 may play roles in the CNS through the PBM-PDZ interactions with the nine PDZ-containing proteins with neuronal functions found to interact with its PBM. Interestingly, SARS-CoV-2 affects multiple organ systems including the CNS. Most patients with COVID-19, caused by SARS-CoV-2, exhibit neurological symptoms. Neurons can be a target of SARS-CoV-2 infection, with localized ischemia in the brain and cell death, highlighting SARS-CoV-2 neurotropism and neuronal infection can be prevented by blocking ACE2 with antibodies [44]. Indeed, the ACE2 receptor is expressed in the brain by endothelial, neuronal and glial cells. After the spreading of virus to CNS, it contacts with ACE2 on neurons, glia, and vascular endothelium [45]. SARS-CoV-2 might also interact directly with sensory neurons, given that sensory dysfunction including cough, olfactory and taste impairments are frequent in infected patients [46,47]. Recently, the loss of the cilia allowing the reception of odor molecules by sensory neurons has been reported after viral infection as well as the presence of viruses in sensory neurons. In addition, the virus invasion in the olfactory bulb as well as the presence of neuroinflammation and viral RNA in several regions of the brain have been shown [47].

Thus, we identified proteins potentially targeted by ACE2 providing new cellular protein partners through PDZ-PBM interactions. This study provides a resource for the research on possible biological pathways involving ACE2. Moreover, SARS-CoV-2 enters the cells through the ACE2 receptor widely expressed, affects organ systems including the CNS and it has therefore pleiotropic effects. Modifications of the interactions between ACE2 and the neuronal proteins identified in our assay could participate to the neurological alterations observed in COVID-19.

## Materials and Methods

### Peptide synthesis

The biotinylated peptide (sequence Biotin-(PEG3)-GFQNTDDVQTSF)(Figure 1C) was synthesized in solid phase using Fmoc strategy (Proteogenix). The sequence encompasses a biotinyl group, a (PEG, polyethylene glycol)_3_ spacer and the C-terminal sequence of the human ACE2 protein (UniProtKB - Q9BYF1) corresponding to the last twelve residues. Peptides were resuspended in the buffer used for holdup.

### Holdup assay

The human PDZ domains library was produced following the high-throughput protocol previously described [13]. Succinctly, the His6-MBP-PDZ constructs were expressed in *E. coli* and all constructs were adjusted to approximately 4 μM concentration in cell lysate soluble fractions using a microcapillary electrophoretic system (Caliper; PerkinElmer, Waltham, MA, USA) and frozen in 96-well plates.

The holdup assay was performed as previously described [11,13] with the biotinylated peptide. Briefly, we measured the fraction of PDZ depletion in the liquid phase during a pull-down experiment. PDZ-containing cell lysates were incubated with the streptavidin resins saturated with biotinylated PBM peptides or with biotin used as a negative control. The supernatant was separated from the resin by a fast filtration step using filter plates. PDZ concentrations were estimated using the microcapillary electrophoretic system.

BI values were calculated using the formula: BI = (Itot-Idepl)/Itot, where Itot is the total intensity of the PDZ peak (biotin control) and Idepl is the intensity of PDZ peak in the peptide depleted reaction. In the holdup buffer, lysozyme was used as internal standard for peak intensity normalization. BI = 0.2 was used as the minimal BI threshold value to define high-confidence PDZ-PBM pairs, as proposed previously [11]. For the detailed protocols, see references [11,13].

We measured peptides interactions against the full human PDZome containing 266 PDZ domains over 272 PDZ domains identified in the human genome [48]. Error bars are standard deviations of the two independent experiments.

### Conversion of BI values to dissociation equilibrium constants

Dissociation constants (Kd) values were calculated from the BIs (Figure 2, Table 1 and Table S1) according to the method published in Gogl *et al*. [15].

BIs were transformed into dissociation constants (K_D_) using the following formula:

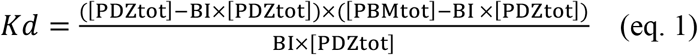

where [PDZ_tot_] and [PBM_tot_] correspond to the total concentrations of the PDZ domain (typically 4 μM) and the PBM peptide, respectively, used during the assay. The PBM_tot_ concentration in the resin during the holdup assay may differ from one peptide to another. Thus, to convert BI values into K_D_ constants, we used the K_D_ values of the three PBM-PDZ interactions previously reported between the ACE2 PBM peptide and three PDZ domains (PDZ1 NHERF3, and the PDZ domains of SHANK1 and SNX27) measured by fluorescence polarization assays [9]. These affinities were used to back-calculate the PBM peptide concentrations in the holdup assays. We have found that all determined BI-*K*_d_ pairs resulted a mean immobilized peptide concentration of 21 μM, a concentration that is coherent with other peptides in the same method [12,14,15]. The minimal BI threshold value of 0.2, defined for significant detectable binders, roughly corresponds to a 80 μM Kd value as previously reported [12].

## Acknowledgements

This research was funded by URGENCE COVID-19 fundraising campaign of Institut Pasteur and the ANR Recherche Action Covid19–FRM PDZCov2 program.

## Conflict of interest

The authors declare no conflict of interest.

## Author contributions

CCS and NW planned the experiments. CCS performed the experiments. CCS and NW analyzed the data. CCS and NW wrote the paper.

## Supplementary Material

Table S1. All BI values obtained by holdup assay using PBM peptide from ACE2 protein.

## Notes

### Competing Interest Statement

The authors have declared no competing interest.

