## Supplemental Table 1 Table S1 for "PDZ-containing proteins targeted by the ACE2 receptor"

**Table S1. All BI values obtained by holdup assay using PBM peptide from ACE2 protein.**

| PDZ name | BI | sd | Kd estimated |
|---|---|---|---|
| SNX27 | 0.87 | 0.03 | 2.57 |
| SHANK3 | 0.75 | 0.03 | 6.08 |
| MAST2 | 0.72 | 0.06 | 6.96 |
| NHERF2 PDZ2 | 0.67 | 0.05 | 9.30 |
| MAST1 | 0.63 | 0.10 | 10.71 |
| SHANK2 | 0.62 | 0.03 | 11.50 |
| PTPN3 | 0.54 | 0.06 | 16.24 |
| SHANK1 | 0.54 | 0.08 | 16.34 |
| NHERF3 PDZ1 | 0.50 | 0.04 | 19.19 |
| NHERF1 PDZ1 | 0.39 | 0.02 | 31.05 |
| PARD3 | 0.28 | 0.05 | 51.12 |
| SHROOM2 PDZ1 | 0.26 | 0.04 | 55.74 |
| SCRIB PDZ3 | 0.23 | 0.00 | 69.12 |
| GRASP | 0.20 | 0.01 | 81.33 |
| MAST3 PDZ1 | 0.20 | 0.02 | |
| RHPN1 PDZ1 | 0.16 | 0.06 | |
| MAGI1 PDZ5 | 0.16 | | |
| NHERF2 PDZ1 | 0.15 | 0.04 | |
| MAGI3 PDZ5 | 0.14 | 0.06 | |
| GRID2IP PDZ2 | 0.14 | 0.21 | |
| PTPN4 PDZ1 | 0.13 | 0.03 | |
| DLG2 PDZ2 | 0.13 | 0.25 | |
| GIPC1 PDZ1 | 0.12 | 0.22 | |
| NHERF PDZ2 | 0.11 | 0.05 | |
| FRMPD4 PDZ1 | 0.11 | 0.02 | |
| PARD3B PDZ1 | 0.10 | 0.01 | |
| PARD3 PDZ3 | 0.09 | 0.07 | |
| DFNB31 PDZ1 | 0.09 | 0.02 | |
| ARHGAP21 PDZ1 | 0.09 | 0.06 | |
| FRMPD3 PDZ1 | 0.07 | 0.05 | |
| MYO18A PDZ1 | 0.06 | 0.02 | |
| MAGI1 PDZ4 | 0.06 | 0.03 | |
| NHERF3 PDZ2 | 0.05 | 0.01 | |

| | | |
|---|---|---|
| PDLIM3 PDZ1 | 0.05 | 0.01 |
| LIN7C PDZ1 | 0.05 | |
| NHERF4 PDZ1 | 0.05 | 0.01 |
| PDLIM1 PDZ1 | 0.04 | 0.07 |
| MAGI1 PDZ6 | 0.04 | |
| ARHGEF12 PDZ1 | 0.04 | 0.00 |
| MPDZ PDZ9 | 0.03 | 0.13 |
| CARD14 PDZ1 | 0.03 | 0.05 |
| ARHGAP23 PDZ1 | 0.03 | 0.09 |
| CNKSR2 PDZ1 | 0.03 | 0.01 |
| CASK PDZ1 | 0.03 | 0.08 |
| SDCBP2 PDZ1 | 0.03 | 0.07 |
| MAGI3 PDZ3 | 0.03 | 0.05 |
| LMO7 PDZ1 | 0.03 | 0.07 |
| PPP1R9A PDZ1 | 0.02 | 0.00 |
| SCRIB PDZ1 | 0.02 | |
| PTPN13 PDZ4 | 0.02 | 0.15 |
| MAGI2 PDZ6 | 0.02 | 0.03 |
| MAGI3 PDZ1 | 0.02 | 0.07 |
| HTRA1 PDZ1 | 0.02 | 0.06 |
| GRIP2 PDZ2 | 0.02 | 0.02 |
| LDB3 PDZ1 | 0.02 | 0.02 |
| PSMD9 PDZ1 | 0.02 | 0.01 |
| InaDl PDZ5 | 0.02 | 0.04 |
| TJP1 PDZ2 | 0.01 | 0.02 |
| PSCDBP PDZ1 | 0.01 | 0.04 |
| LNX2 PDZ4 | 0.01 | 0.06 |
| PDZD11 PDZ1 | 0.01 | 0.02 |
| PPP1R9B PDZ1 | 0.01 | 0.01 |
| CARD11 PDZ1 | 0.01 | 0.03 |
| DVL3 PDZ1 | 0.01 | 0.08 |
| PDZRN3 PDZ2 | 0.01 | |
| InaDl PDZ9 | 0.01 | 0.04 |
| AHNAK1 PDZ1 | 0.00 | |
| CNKSR3 PDZ1 | 0.00 | |
| SIPA1L1 PDZ1 | 0.00 | 0.03 |
| MPDZ PDZ7 | 0.00 | 0.02 |
| RADIL PDZ1 | 0.00 | 0.06 |
| PDZD7 PDZ1 | 0.00 | 0.06 |
| MAST4 PDZ1 | 0.00 | 0.04 |
| GRIP2 PDZ5 | 0.00 | 0.03 |
| SNTG1 PDZ1 | -0.01 | 0.02 |
| PDZD2 PDZ4 | -0.01 | 0.01 |

| | | |
|---|---|---|
| InaDl PDZ2 | -0.01 | 0.01 |
| GRIP1 PDZ3 | -0.01 | |
| DEPTOR PDZ1 | -0.01 | |
| InaDl PDZ7 | -0.01 | 0.08 |
| MPDZ PDZ4 | -0.01 | 0.02 |
| DLG5 PDZ4 | -0.01 | 0.04 |
| LNX2 PDZ2 | -0.01 | 0.02 |
| GRIP1 PDZ1 | -0.01 | 0.10 |
| SNTB1 PDZ1 | -0.01 | 0.10 |
| InaDl PDZ4 | -0.01 | 0.04 |
| PDZRN3 PDZ1 | -0.01 | 0.13 |
| PARD3B PDZ3 | -0.01 | 0.02 |
| LIN7B PDZ1 | -0.01 | 0.05 |
| SDCBP2 PDZ2 | -0.01 | 0.04 |
| DLG2 PDZ1 | -0.01 | 0.02 |
| NHERF3 PDZ3 | -0.01 | 0.02 |
| TJP1 PDZ1 | -0.01 | 0.05 |
| MLLT4 PDZ1 | -0.01 | 0.01 |
| DLG3 PDZ3 | -0.02 | 0.01 |
| LAP2 PDZ1 | -0.02 | 0.09 |
| RIMS2 PDZ1 | -0.02 | 0.11 |
| RGS12 PDZ1 | -0.02 | 0.03 |
| DLG4 PDZ2 | -0.02 | 0.01 |
| SCRIB PDZ4 | -0.02 | 0.02 |
| PARD3 PDZ2 | -0.02 | 0.02 |
| SHROOM4 PDZ1 | -0.02 | 0.07 |
| PTPN13 PDZ3 | -0.02 | 0.02 |
| MPP4 PDZ1 | -0.02 | 0.04 |
| PTPN13 PDZ1 | -0.02 | 0.02 |
| NHERF4 PDZ3 | -0.02 | 0.03 |
| RIMS1 PDZ1 | -0.02 | 0.07 |
| PDZRN4 PDZ2 | -0.02 | 0.00 |
| NOS1 PDZ1 | -0.02 | 0.00 |
| FRMPD2 PDZ1 | -0.02 | 0.05 |
| LNX1 PDZ4 | -0.02 | 0.06 |
| PDZD9 PDZ1 | -0.02 | 0.11 |
| GIPC3 PDZ1 | -0.02 | 0.05 |
| DLG5 PDZ2 | -0.03 | 0.02 |
| STXB4 PDZ1 | -0.03 | 0.07 |
| GRID2IP PDZ1 | -0.03 | 0.02 |
| SIPA1L2 PDZ1 | -0.03 | 0.00 |
| SCRIB PDZ2 | -0.03 | 0.06 |
| PRX PDZ1 | -0.03 | 0.01 |

| | | |
|---|---|---|
| PDZD2 PDZ2 | -0.03 | 0.02 |
| FRMPD1 PDZ1 | -0.03 | 0.03 |
| InaDl PDZ6 | -0.03 | 0.00 |
| MAGI1 PDZ1 | -0.03 | 0.06 |
| PDZD7 PDZ2 | -0.03 | 0.03 |
| PDZD2 PDZ3 | -0.03 | 0.02 |
| MPP5 PDZ1 | -0.03 | 0.01 |
| SNTA1 PDZ1 | -0.03 | 0.03 |
| MAGI2 PDZ2 | -0.03 | 0.01 |
| PDZD2 PDZ6 | -0.03 | 0.05 |
| PICK1 PDZ1 | -0.03 | |
| DLG4 PDZ3 | -0.03 | 0.05 |
| APBA2 PDZ1 | -0.03 | |
| FRMPD2 PDZ3 | -0.03 | 0.03 |
| GORASP2 PDZ1 | -0.03 | 0.04 |
| MAGI3 PDZ6 | -0.03 | 0.01 |
| MPDZ PDZ6 | -0.03 | 0.03 |
| MPDZ PDZ13 | -0.03 | 0.03 |
| MAGI1 PDZ2 | -0.04 | 0.00 |
| APBA1 PDZ2 | -0.04 | |
| MAGI2 PDZ5 | -0.04 | 0.02 |
| RGS3 PDZ1 | -0.04 | 0.07 |
| DLG3 PDZ1 | -0.04 | 0.06 |
| PDLIM4 PDZ1 | -0.04 | 0.00 |
| MAGI2 PDZ1 | -0.04 | 0.01 |
| MAGI1 PDZ3 | -0.04 | 0.01 |
| MPP3 PDZ1 | -0.04 | 0.02 |
| IL16 PDZ3 | -0.04 | 0.01 |
| GRIP2 PDZ7 | -0.04 | 0.06 |
| AHNAK2 PDZ1 | -0.04 | |
| SIPA1 PDZ1 | -0.04 | 0.01 |
| MAGI2 PDZ4 | -0.04 | 0.01 |
| LIN7A PDZ1 | -0.04 | 0.05 |
| PDZD8 PDZ1 | -0.04 | 0.00 |
| DLG2 PDZ3 | -0.04 | 0.05 |
| GRIP2 PDZ3 | -0.04 | 0.00 |
| MPP7 PDZ1 | -0.04 | 0.00 |
| GRIP2 PDZ6 | -0.04 | 0.02 |
| GRIP1 PDZ7 | -0.04 | 0.02 |
| SHROOM3 PDZ1 | -0.04 | 0.03 |
| MPDZ PDZ3 | -0.04 | 0.07 |
| GIPC2 PDZ1 | -0.05 | 0.04 |
| PREX2 PDZ2 | -0.05 | 0.01 |

| | | |
|---|---|---|
| MPDZ PDZ2 | -0.05 | 0.03 |
| PREX1 PDZ2 | -0.05 | 0.00 |
| MAGI3 PDZ2 | -0.05 | 0.00 |
| PDZRN4 PDZ1 | -0.05 | 0.06 |
| MPDZ PDZ1 2 | -0.05 | 0.01 |
| PCLO PDZ1 | -0.05 | 0.01 |
| MPP6 PDZ1 | -0.05 | 0.00 |
| PDLIM7 PDZ1 | -0.05 | 0.00 |
| SYNPO2L PDZ1 | -0.05 | 0.04 |
| PTPN13 PDZ5 | -0.05 | 0.04 |
| SNTG2 PDZ1 | -0.05 | 0.02 |
| SYNP2 PDZ1 | -0.05 | 0.02 |
| MPDZ PDZ11 | -0.05 | 0.04 |
| HTRA4 PDZ1 | -0.05 | 0.01 |
| MAGI3 PDZ4 | -0.05 | 0.01 |
| DFNB31 PDZ2 | -0.05 | 0.02 |
| PDLIM2 PDZ1 | -0.05 | 0.00 |
| SIPA1L3 PDZ1 | -0.05 | 0.02 |
| MPDZ PDZ5 | -0.05 | 0.04 |
| PDZD4 PDZ1 | -0.05 | 0.02 |
| PARD6G PDZ1 | -0.05 | 0.00 |
| MPDZ PDZ8 | -0.06 | 0.03 |
| HTRA3 PDZ1 | -0.06 | 0.00 |
| IL16 PDZ1 | -0.06 | 0.03 |
| PDZD7 PDZ3 | -0.06 | 0.05 |
| RAPGEF2 PDZ1 | -0.06 | |
| MAGI2 PDZ3 | -0.06 | 0.01 |
| LRRC7 PDZ1 | -0.06 | 0.01 |
| GORASP1 PDZ1 | -0.06 | 0.06 |
| DLG5 PDZ3 | -0.06 | 0.08 |
| SYNJ2BP PDZ1 | -0.06 | 0.02 |
| InaDl PDZ3 | -0.06 | 0.05 |
| APBA2 PDZ2 | -0.06 | 0.01 |
| RAPGEF6 PDZ1 | -0.06 | 0.02 |
| NHERF3 PDZ4 | -0.06 | 0.19 |
| MAGIX PDZ1 | -0.06 | 0.01 |
| NHERF4 PDZ2 | -0.06 | 0.10 |
| LNX1 PDZ1 | -0.07 | 0.04 |
| DLG1 PDZ2 | -0.07 | 0.03 |
| MPDZ PDZ1 0 | -0.07 | 0.01 |
| LNX1 PDZ2 | -0.07 | |
| PARD3B PDZ2 | -0.07 | 0.06 |
| LNX1 PDZ3 | -0.07 | 0.03 |

| | | |
|---|---|---|
| TJP2 PDZ1 | -0.07 | 0.03 |
| GRIP1 PDZ2 | -0.07 | 0.02 |
| GRIP1 PDZ5 | -0.07 | 0.03 |
| LNX2 PDZ1 | -0.07 | 0.01 |
| INTU PDZ1 | -0.07 | 0.05 |
| MPDZ PDZ1 | -0.07 | 0.01 |
| MPP1 PDZ1 | -0.07 | 0.05 |
| FRMPD2 PDZ2 | -0.07 | 0.03 |
| DLG1 PDZ3 | -0.07 | 0.02 |
| PDLIM5 PDZ1 | -0.08 | 0.02 |
| SDCBP PDZ1 | -0.08 | 0.02 |
| HTRA2 PDZ1 | -0.08 | 0.01 |
| TIAM1 PDZ1 | -0.08 | 0.04 |
| APBA1 PDZ1 | -0.08 | |
| RHPN2 PDZ1 | -0.08 | 0.04 |
| TJP3 PDZ2 | -0.08 | 0.08 |
| GRIP2 PDZ4 | -0.08 | 0.09 |
| SNTB2 PDZ1 | -0.08 | 0.07 |
| PARD6A PDZ1 | -0.08 | 0.00 |
| TJP3 PDZ3 | -0.08 | 0.01 |
| SDCBP PDZ2 | -0.09 | 0.04 |
| InaDl PDZ10 | -0.09 | 0.04 |
| IL16 PDZ4 | -0.09 | 0.00 |
| PTPN13 PDZ2 | -0.09 | 0.10 |
| GOPC PDZ1 | -0.09 | 0.02 |
| TX1B3 PDZ1 | -0.09 | 0.04 |
| PDZD2 PDZ1 | -0.09 | 0.02 |
| IL16 PDZ2 | -0.09 | 0.03 |
| APBA3 PDZ2 | -0.10 | |
| LIMK1 PDZ1 | -0.10 | 0.08 |
| GRIP1 PDZ4 | -0.10 | 0.06 |
| DLG5 PDZ1 | -0.10 | |
| PREX1 PDZ1 | -0.10 | 0.05 |
| PARD6B PDZ1 | -0.10 | 0.01 |
| TJP3 PDZ1 | -0.10 | 0.02 |
| TJP2 PDZ3 | -0.10 | |
| APBA3 PDZ1 | -0.11 | 0.18 |
| LIMK2 PDZ1 | -0.11 | 0.03 |
| GRIP2 PDZ1 | -0.11 | 0.13 |
| USH1C PDZ3 | -0.11 | 0.01 |
| PREX2 PDZ1 | -0.11 | 0.06 |
| NHERF4 PDZ4 | -0.11 | 0.08 |
| MPP2 PDZ1 | -0.12 | 0.02 |

| | | |
|---|---|---|
| TJP2 PDZ2 | -0.12 | 0.04 |
| TIAM2 PDZ1 | -0.12 | 0.13 |
| PDZD2 PDZ5 | -0.13 | 0.02 |
| USH1C PDZ1 | -0.13 | |
| DVL1 PDZ1 | -0.14 | 0.18 |
| DLG1 PDZ1 | -0.14 | 0.19 |
| TJP1 PDZ3 | -0.14 | 0.10 |
| ARHGEF11 PDZ1 | -0.18 | 0.19 |
| InaDl PDZ8 | -0.24 | 0.29 |
| DFNB31 PDZ3 | -0.24 | 0.33 |
| USH1C PDZ2 | -0.25 | 0.10 |
| DLG3 PDZ2 | -0.27 | 0.38 |
| GRIP1 PDZ6 | -0.27 | 0.36 |
| CNKSR1 PDZ1 | -0.31 | 0.38 |
| InaDl PDZ1 | -0.34 | 0.31 |
| DVL2 PDZ1 | -0.38 | 0.57 |
| DLG4 PDZ1 | -0.50 | |
| LNX2 PDZ3 | | |
